# Dietary Menaquinone-9 Supplementation Does Not Influence Bone Tissue Quality or Bone Mineral Density in Mice

**DOI:** 10.1101/2025.01.29.635567

**Authors:** Minying Liu, Chongshan Liu, Nicolas Cevallos, Benjamin N Orbach, Christopher J Hernandez, Xueyan Fu, Jennifer Lee, Sarah L Booth, M Kyla Shea

**Affiliations:** Human Nutrition Research Center on Aging, Tufts University, Boston, MA; Orthopaedic Surgery, University of California, San Francisco, CA; Sibley School of Mechanical and Aerospace Engineering, Cornell University, Ithaca, NY; Graduate School of Biomedical Sciences, Tufts University, Boston, MA

**Keywords:** skeletal health, bone tissue quality, bone mineral density, vitamin K, menaquinone

## Abstract

Vitamin K has been implicated in skeletal health because vitamin K-dependent proteins are present in bone. While there are multiple forms of vitamin K, most research has focused on phylloquinone, which is found mainly in plant-based foods, and its metabolite menaquinone-4 (MK4). However, there are additional forms of vitamin K that are bacterially produced that appear to influence bone health but have not yet been studied extensively. Herein, we evaluated the effects of menaquinone-9 (MK9), a bacterially produced form of vitamin K on bone tissue quality and density in young mice. Four-week-old male (n=32) and female (n=32) C57BL/6 mice were supplemented with 0.06 mg/kg diet or 2.1 mg/kg diet of MK9 for 12 weeks. During week 11, a sub-group of mice (n=7/sex/group) received daily deuterium-labeled MK9 to trace its metabolic fate in bone. Liver MK4 and MK9 were significantly higher in mice fed 2.1 mg MK9/kg compared to those receiving 0.06 mg MK9/kg, regardless of sex (all p ≤ 0.017). MK4 was the only vitamin K form detected in bone, with 63-67% of skeletal MK4 in mice fed 2.1 mg MK9/kg derived from deuterium-labeled MK9. Femoral tissue strength, maximum bending moment, section modulus, and bone mineral density did not differ significantly across diet groups in either sex (all p≥0.083). Cross-sectional area (p=0.003) and moment of inertia (p=0.001) were lower in female mice receiving 2.1 mg MK9/kg compared to those receiving 0.06 mg MK9/kg, but no differences were found in male mice. Higher bone MK4 concentrations did not correlate with higher bone tissue quality or density. Despite dietary MK9 being a dietary precursor to MK4 in bone, dietary MK9 supplementation did not affect bone tissue quality or bone mineral density.

**Lay summary:** Most research about vitamin K and bone health has focused on phylloquinone, the plant-based vitamin K form, and its metabolite menaquinone-4. Because interest in bacterially produced forms of vitamin K, which are abundant in the intestinal microbiome, is growing, we evaluated the effect of menaquinone-9 (a bacterially-produced form of vitamin K) on skeletal health. We supplemented mice with low and high doses of menaquinone-9 and also used stable-isotope labeled menaquinone-9 to trace its conversion to menaquinone-4 in bone. We found menaquinone-9 served as a precursor to menaquinone-4 in bone, but menaquinone-9 supplementation did not improve bone health.

## INTRODUCTION

Vitamin K, an essential fat-soluble nutrient, has been implicated in bone health (1) based on its role as an enzymatic cofactor required for the post-translational carboxylation of certain proteins (vitamin K-dependent proteins) (2). One such protein is osteocalcin, the predominate non-collagenous protein in bone. Once carboxylated, osteocalcin binds to calcium in the bone matrix (3). There are multiple naturally-occurring forms of vitamin K which are capable of carboxylation. The plant-based form, phylloquinone (PK) is found in green leafy vegetables and vegetable oils (4). Menaquinones (MKs) are a class of vitamin K compounds that are primarily bacterially produced and are therefore abundant in the intestinal microbiome. At least 13 MK forms have been identified that differ structurally from PK in the length and saturation of their prenylated side chain, with the number of prenyl units differentiating among the specific MKs (MKn) (5, 6).

Among the menaquinones, MK4 is unique because it is not known to be made by bacteria. Instead, it is synthesized in several extra-hepatic tissues from other forms of vitamin K, including MKs that are bacterially synthesized (7–9). It was recently demonstrated that MK4 is found in bone (9), but the role of dietary vitamin K as a precursor to MK4 in bone has not yet been fully elucidated. This is an important gap to address because MK4 signaling pathways have been implicated in bone tissue homeostasis (10, 11).

Most observational studies and randomized clinical trials evaluating the role of vitamin K in skeletal health have focused primarily on PK and MK4, with age-related bone loss measured using bone mineral density (BMD) as the main outcome. However, these studies have yielded inconsistent results (12). A systematic review and meta-analysis of randomized controlled trials that tested the effect of PK or MK4 supplementation on age-related bone loss concluded there is little evidence that vitamin K affects BMD or vertebral fractures. The authors reported a potentially beneficial effect of PK or MK4 supplementation on clinical fracture risk, but concluded additional studies are needed to confirm this (13). While fragility fractures are often associated with low BMD, they also occur in individuals with normal BMD (14). Bone tissue quality—encompassing bone strength and mechanical properties—is another critical aspect of bone health not captured by BMD (15–18). An impairment in bone tissue quality can reduce whole bone strength without altering BMD (19). In mice, knockout of the vitamin K-dependent protein osteocalcin results in impaired bone tissue composition and quality with only minor changes in bone geometry or BMD (20, 21). Hence, vitamin K-induced alteration of bone tissue quality may not be reflected in BMD or other indices of bone geometry and mass. Furthermore, there are limited data on the potential effects of bacterially-produced MKs on bone health outcomes. Addressing this gap is essential for understanding the broader role of vitamin K in skeletal health.

To clarify the role of bacterially-produced MK forms in bone tissue, we supplemented male and female C57BL/6 mice with two different doses of menaquinone-9 (MK9) in their diet for 12 weeks. We chose MK9 because: (I) MK9 is a MK form produced by bacteria. (II) Of the MK forms currently characterized in the food supply, MK9 is the most abundant, especially in dairy and fermented products (6). (III) Through use of stable isotopes, we have already characterized the metabolic fate of dietary MK9 in C57BL/6 mice, including the conversion of MK9 to MK4 in extra-hepatic tissues other than bone (9), which serves as a quality control from which to compare our findings. We also evaluated the conversion of dietary MK9 to MK4 in skeletal tissue using stable isotopes. We hypothesized that dietary supplementation with MK9 would positively influence bone tissue quality and density and serve as a precursor to MK4 in bone tissue.

## MATERIALS AND METHODS

### Study design and experimental diets

Three-week-old male (n=32) and female (n=32) C57BL/6 mice were acquired from Charles River Laboratories and housed in single cages at the Tufts University Human Nutrition Research Center on Aging (HNRCA) Comparative Biology Unit (CBU). Upon arrival, mice were acclimated to an AIN-93G diet (Inotiv, TD.94045), which contained 1mg/kg PK per manufacture, for one week. At 4 weeks of age, mice were divided at random into 2 experimental groups to receive diets containing varying amounts of MK9 for 12 weeks (22). Males and females were randomized separately to one of the two experimental groups (**Figure 1**): (I) diet containing 2.1 mg/kg MK9, (II) diet containing 0.06 mg/kg MK9. Diets were composed of a mixture of the vitamin K-free basal diet (Inotiv, TD.120060) with two different amounts of MK9. An MK9 content of 2.1 mg/kg corresponds to the recommended vitamin K intake, which was based on the molar equivalent of MK9 to the currently recommended concentration of vitamin K for rodents (1mg/kg PK), following a protocol developed by the Vitamin K Laboratory at the HNRCA (23). In contrast, an MK9 content of 0.06 mg/kg indicates a vitamin K intake below the recommended level, which was molar equivalent to 31 ± 0.45 μg PK/kg contained in vitamin K-free basal diet (9, 24). The Vitamin K laboratory previously demonstrated that feeding mice a diet containing MK9 (as the only form of vitamin K in the diet) in an equal molar amount to the current rodent diet recommendations showed no sign of vitamin K-deficiency (9, 24). Mice were pair-fed between diet groups to minimize the possibility that any outcome differences would be due to differences in the total amount of food consumed.

**Figure 1.**
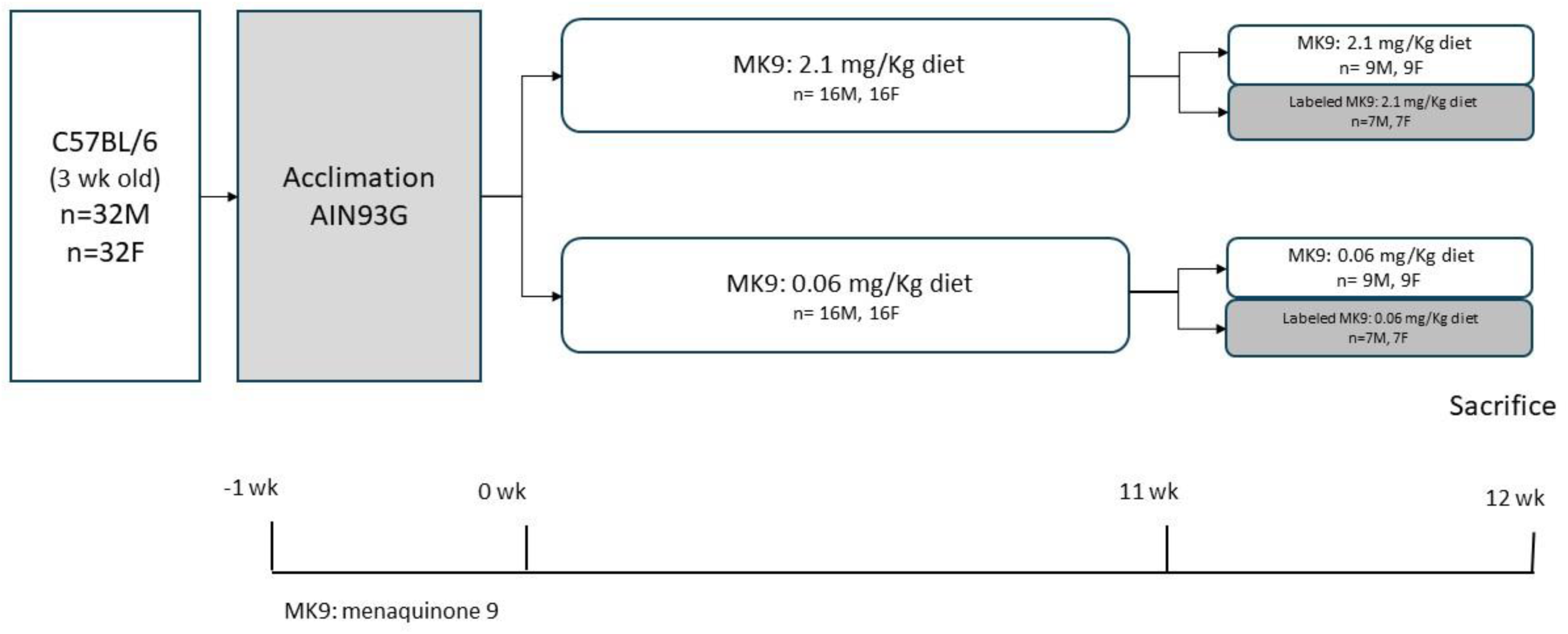
Animal Study Design. Four-week-old male (n=32) and female (n=32) C57BL/6 mice were supplemented with 0.06 mg/kg and 2.1 mg/kg of menaquinone-9 (MK9) for 12 weeks. During the final week, a sub-group of mice (n=7/sex/group) received deuterium-labeled MK9 to trace its metabolic fate in bone.

To determine whether MK9 from the diet serves as a precursor to MK4 present in bone a subset of mice (n=14, 7 males and 7 females) from each group were randomly selected to receive daily diets containing deuterium-labeled MK9 (^2^H_7_MK9) (in the same amount as in the initial diet, mixed with vitamin K-free basal diet using the protocol developed by the Vitamin K Laboratory) during week 11 of the experiment (the last week prior to sacrifice) (**Figure 1**). MK9 concentrations in study diets were measured by liquid chromatography-mass spectrometry (LC-MS)(25), and diet MK9 concentrations were 0.06±0.004 mg MK9/kg, 0.07±0.006 mg ^2^H_7_MK9/kg, 2.03±0.12 mg MK9/kg, and 2.47±0.18 mg ^2^H_7_MK9/kg, respectively.

Animals were euthanized at 16 weeks of age. After sacrifice, mice carcasses were scanned for bone mineral density measurement, and then femurs, tibias, and liver were collected immediately after euthanasia and stored and maintained at -80 ℃ for bone biomechanical testing and vitamin K analyses. All animal experiments and procedures were approved by the Institutional Animal Care and Use Committee at the HNRCA at Tufts University.

### Bone tissue quality

Right femurs were dissected and shipped to the Hernandez Laboratory at the University of California, San Francisco, where femur length, cross-sectional geometry, and maximum bending force were measured, as described elsewhere (22, 26). In brief, femur length was measured using digital calipers from the greater trochanter to the lateral condyle. The femoral diaphyseal cross-section was imaged with a micro-computed tomography scanner at a voxel size of 10 μm to assess femoral cross-sectional geometry. The measurements included cortical cross-sectional area, moment of inertia about the medial-lateral axis (I), the distance from the neutral axis to bone surface (c), and section modulus (I/c). For mechanical testing, femurs were thawed to room temperature and kept hydrated. They were then subjected to three-point bending tests in the anterior-posterior direction using a servo-hydraulic testing device (858 Mini Bionix; MTS) until failure occurred (loading rate = 0.1 mm/s, span length = 8 mm between outer loading pins). The maximum bending force and displacement were recorded. The maximum bending force was used to calculate maximum bending moment (max force x span length/4). Bone tissue strength was calculated using maximum bending moment and cross-sectional geometric measurements (maximum bending moment / section modulus).

### Bone mineral density

After sacrifice, mice carcasses were scanned using the InAlyzer2 dual-energy X-ray absorptiometry (DEXA) system, Model M (Medikors Inc., Korea) at HNRCA, with details described elsewhere (27). In brief, a quality control phantom was scanned prior to each measurement. Each carcass was positioned in the center of the scanning area in a prone position, with its arms and legs extended outward and its tail aligned straight behind the body. Scans were performed with a pixel size of 103×106 μm and a pixel pitch of 48 μm (108 μm in analysis). Total body BMD (g/cm²) was obtained using InAlyzer software version 3.2.3.

### Vitamin K analyses

Measurements of MKs in bone and liver were conducted in the Vitamin K Laboratory at HNRCA as previously described (25). Briefly, liver tissues (0.15 g) were homogenized in phosphate-buffered saline (PBS) by use of a Powergen homogenizer (Fisher Scientific). Left and right tibias (in total of 0.1g) collected at sacrifice were fragmented using a stainless-steel tissue pulverizer (Biospec) with the addition of liquid nitrogen to ensure thorough pulverization. The resulting bone powder was transferred into 2 mL tubes pre-filled with beads, followed by the addition of 1 mL PBS. Samples were then homogenized using an Omni Bead Ruptor 24 Homogenizer, running two cycles of 1 minute each. Homogenized tissue samples were extracted using hexane, further purified using solid-phase extraction, and then quantified using Quadrupole Time-of-Flight Mass Spectrometry (Q-TOF-MS). To differentiate deuterium-labeled vitamin K forms, a ProntoSil C30 column is required for the separation of stereoisomers (28).

### Statistical power and analyses

Sample size calculations were based on the work of Guss et al (22), who reported a decrease of 3.6 N/mm^2^ in femoral peak moment bending/I/c in 4-week-old C57BL/6 male mice treated with oral antibiotics for 12 weeks compared to mice that received no antibiotic treatment (p<0.05). The between group standard deviation was 2.5 (22). Fifteen mice per group provided 92% power to detect a between group difference of 3.6, using a two-tailed alpha of 0.05. With 2 experimental groups, we had at least 85% power to detect differences comparable to 3.6 N/mm^2^ in Peak Moment in bending/I/c, accounting for up to 6 pair-wise comparisons.

Data are presented as means ± SD. Statistical analyses were performed using RStudio (v.4.3.1, 2023). The Shapiro-Wilk test was used to assess normality, and the Levene’s test was employed to evaluate equal variance. For outcome measures that satisfied parametric assumptions, an independent two-sample t-test was conducted to compare femoral tissue strength (primary outcome), other skeletal outcomes, liver, and bone vitamin K concentrations between diet groups. When parametric assumptions were not met, data were log-transformed to achieve normality, and parametric assumptions were reassessed. The correlations between bone MK4 concentrations and bone biomechanical and geometry outcomes were assessed using the Pearson correlation coefficient. Analyses were conducted separately for males and females, with statistical significance set at p < 0.05. Normalization by body weight was achieved by dividing bone biomechanical measurements by body weight.

## RESULTS

Eight mice died prior to the end of the study (7 males and 1 female). The highest mortality rate was observed among male mice receiving 0.06 mg/kg MK9. In contrast, all female mice receiving 0.06 mg/kg MK9 treatment survived. Additionally, one mouse of each sex was lost among those receiving 2.1 mg/kg MK9 (See Supplemental **Figure S1**).

### Bone tissue quality and density

Femoral tissue strength was not different across diet groups for either sex (male p=0.725; female p=0.302, **Table 1**). Similarly, there were no differences in maximum bending moment between diet groups for either male or female mice (p=0.763 and 0.083 respectively, **Table 1**) or in sectional modulus in male or female mice between the diet groups (p=0.873 and 0.277 respectively, **Table 1**).

**Table 1.** Femoral biomechanics, geometry and total body bone mineral density measurements of male and female mice receiving diets supplemented with MK9.

|  | Male |  | Female |  |
| --- | --- | --- | --- | --- |
|  | 0.06mg/kg (n=7) | 2.1mg/kg (n=11) | 0.06mg/kg (n=13) | 2.1mg/kg (n=11) |
| Tissue strength (Mpa) | 153.3±13.6 | 156±16.3 | 171.8±16.7 | 164.6±14.8 |
| *P values* | 0.725 |  | 0.302 |  |
| Maximum moment (N*mm) | 33.3±2.4 | 33.7±3.5 | 37.9±3 | 35.4±3.7 |
| *P values* | 0.763 |  | 0.083 |  |
| Section modulus (mm<sup>3</sup>) | 0.22±0.02 | 0.22±0.01 | 0.22±0.02 | 0.21±0.02 |
| *P values* | 0.873 |  | 0.277 |  |
| Moment of inertia (mm<sup>4</sup>) | 0.13±0.01 | 0.13±0.01 | 0.14±0.01 | 0.13±0.01 |
| *P values* | 0.46 |  | 0.003 |  |
| Cross sectional area (mm<sup>2</sup>) | 0.86±0.07 | 0.9±0.04 | 0.95±0.07 | 0.87±0.03 |
| *P values* | 0.205 |  | 0.001 |  |
| Femur length (mm) | 15.6±0.3 | 15.6±0.5 | 15.6±0.4 | 15.8±0.3 |
| *P values* | 0.774 |  | 0.288 |  |
| Bone mineral density (g/cm<sup>2</sup>) | 0.06±0.004 | 0.06±0.004 | 0.06±0.002 | 0.06±0.003 |
| *P values* | 0.306 |  | 0.947 |  |

Values are mean±SD. Differences between diet groups were determined using parametric unpaired two sample t-test. Males and female were analyzed separately with statistical significance set as p<0.05. Total body bone mineral density is presented, with all other measurements analyzed using femurs.

Female mice receiving 0.06 mg/kg MK9 had higher moment of inertia and cross-sectional area compared to female mice receiving 2.1mg/kg MK9 (all p-values≤0.003). No significant differences in these two measurements were observed among the male mice across diet groups (all p-values≥0.205, **Table 1**). No significant differences were observed in femur length across diet groups for either male or female mice (p-values≥0.288, **Table 1**). In addition, total body bone mineral density did not differ between diet groups in either sex (all p-values≥0.306, **Table 1**).

Body weight did not differ between diet groups for either male or female mice (all p-values≥0.345, see Supplemental **Figure S2**). When normalized to body weight, femoral tissue strength, maximum moment, section modulus, moment of inertia, and femur length in both male and female mice remained unchanged (see Supplemental **Table S1**). The difference in femoral cross-sectional area for female mice was not statistically significant (p=0.060).

### Tissue Vitamin K Content

We detected MK4 and MK9 in the liver, with significantly higher concentrations of both observed in mice fed 2.1 mg MK9/kg compared to those receiving 0.06 mg MK9/kg, regardless of sex (all p ≤ 0.017, **Figure 2A–D**). Furthermore, liver concentrations of MK9 were higher than those of MK4. MK4 was the only vitamin K form detected in bone. Mice receiving 0.06 mg/kg MK9 had significantly lower bone MK4 concentrations compared to mice receiving 2.1mg/kg MK9 in both sexes (all p values<0.001; **Figure 3 AB**). In the subset of mice fed deuterium-labeled MK9 for the week prior to sacrifice, deuterium-labeled MK4 was detected in bone tissue of both male and female mice that received 2.1 mg/kg labeled MK9 (mean ± SD at 22 ± 9 pmol/g for males and 37 ± 21 pmol/g for females, **Table 2**). In addition, 63-67% of MK4 measured in bone was derived from labeled MK9 (**Table 2**). In female mice, bone tissue MK4 concentrations were significantly inversely correlated with cross-sectional area (Pearson r = - 0.55, p = 0.008) and nearly significantly inversely correlated with moment of inertia (Pearson r = -0.40, p = 0.066). Otherwise, bone tissue MK4 concentrations did not correlate with any other bone outcome analyzed (Pearson r coefficients range -0.25 to +0.29, all p values ≥ 0.25; **Figure S3**), and bone MK4 concentrations were not correlated with any bone outcome in male mice (Pearson r range + 0.01 to + 0.30, all p ≥ 0.35; **Figure S3**).

**Figure 2.**
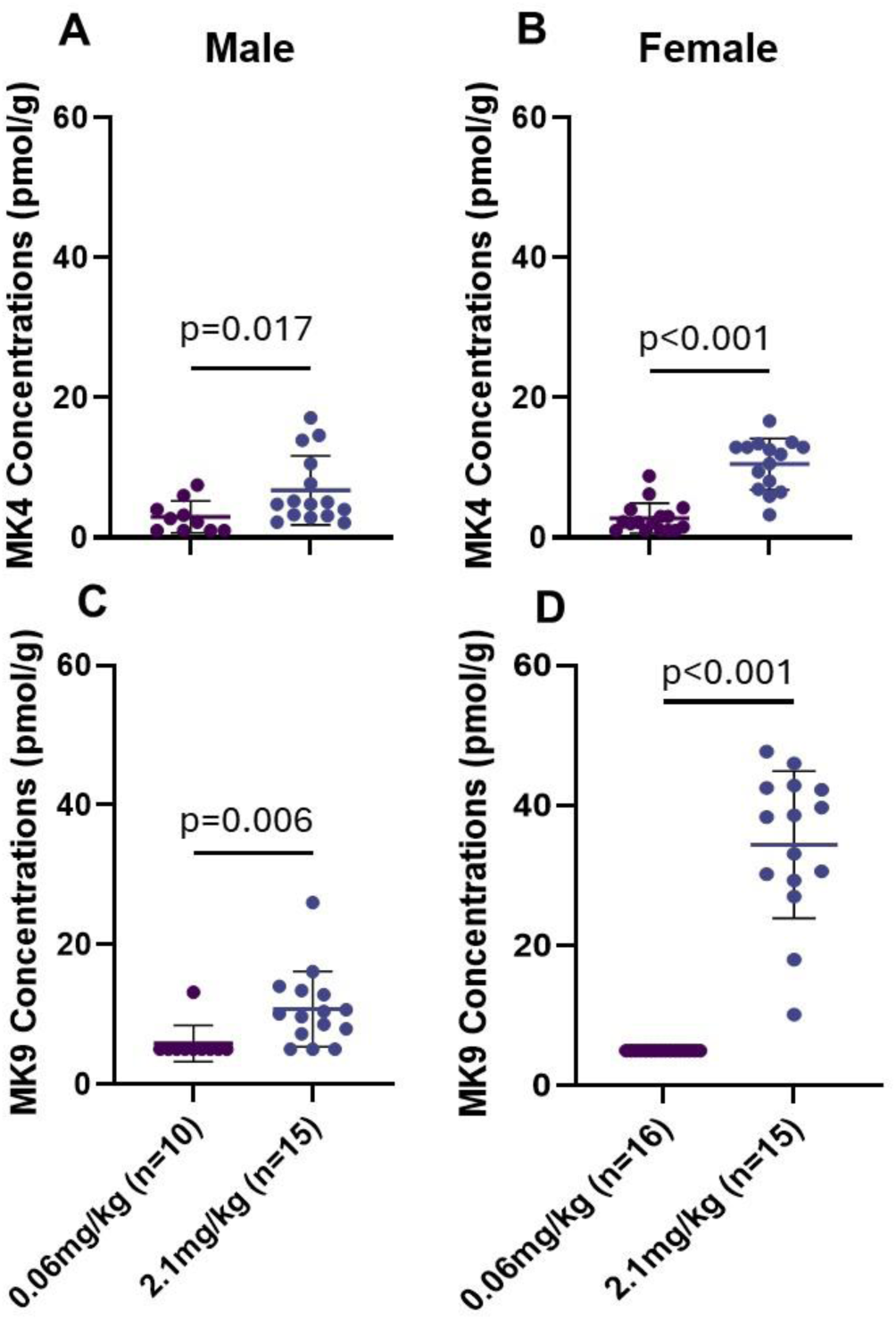
Liver MK4 (A, B) and MK9 (C, D) concentrations (pmol/g) of male (A, C) and female (B, D) mice receiving diets supplemented with MK9. Differences between diet groups were determined using parametric unpaired two sample t-test. Males and female were analyzed separately with statistical significance set as p<0.05. Data are presented as mean ±SD.

**Figure 3.**
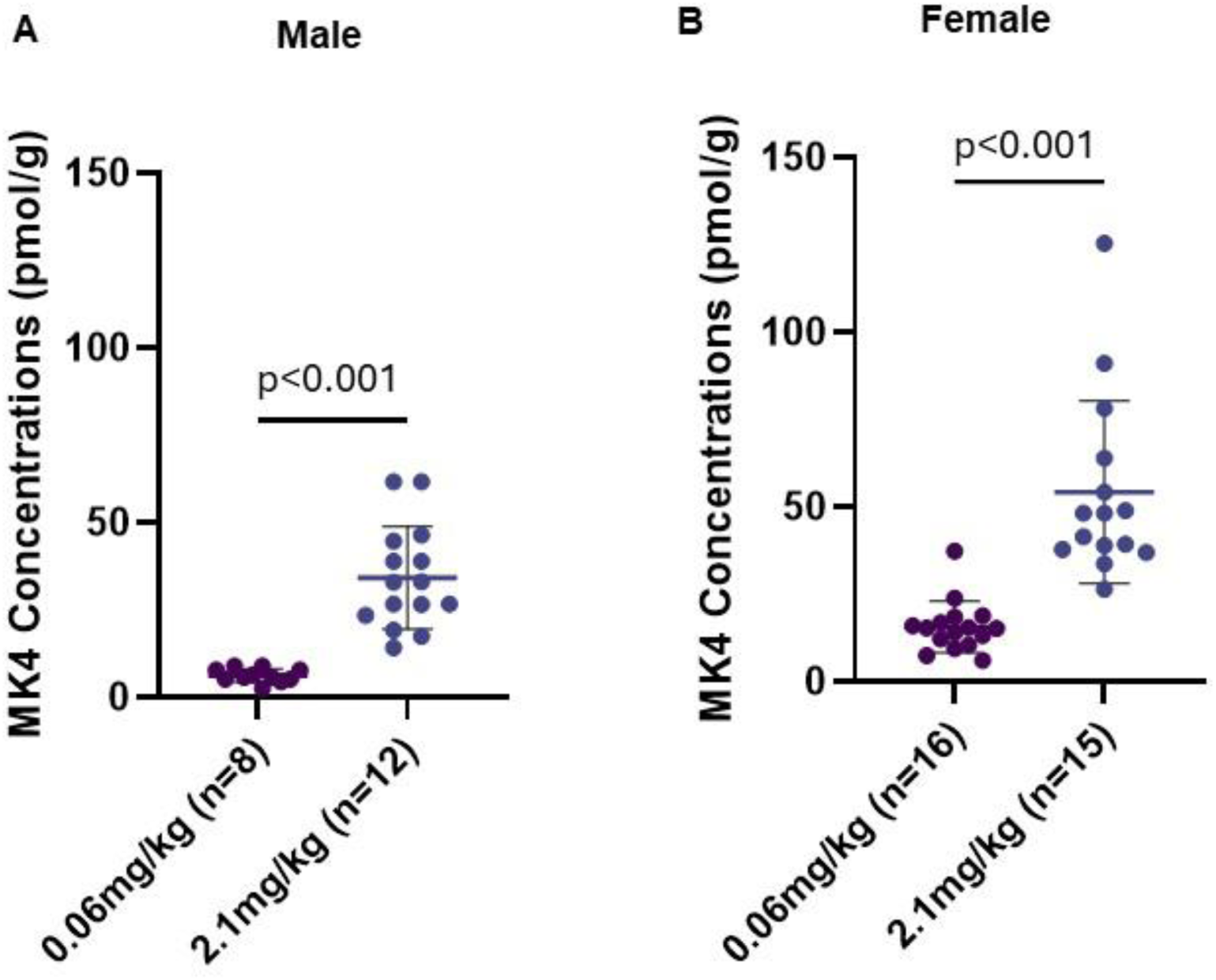
Bone MK4 concentrations (pmol/g) of male (A) and female (B) mice receiving diets supplemented with MK9. Differences between diet groups were determined using parametric unpaired two sample t-test. Males and female were analyzed separately with statistical significance set as p<0.05. Data are presented as mean ±SD.

**Table 2.** Bone MK4 concentrations of male and female mice receiving diets supplemented with deuterium-labeled MK9.

| VK form | Male |  |  | Female |  |  |
| --- | --- | --- | --- | --- | --- | --- |
|  | 0.06mg/kg (n=4) | 2.1mg/kg (n=6) | *P values* | 0.06mg/kg (n=7) | 2.1mg/kg (n=7) | *P values* |
| Unlabeled MK4 (pmol/g) | 4 ± 2 | 11 ± 3 | 0.009 | 12 ± 5 | 23 ± 15 | 0.066 |
| Labeled MK4 (pmol/g) | ND | 22 ± 9 | 0.002 | ND | 37 ± 21 | <0.001 |
| Total MK4 (pmol/g) | 5 ± 2 | 32 ± 12 | 0.002 | 13 ± 5 | 60 ± 35 | <0.001 |
| % of labeled MK4 (of total VK) | NC | 66.6 ± 3.4 | <0.001 | NC | 62.6 ± 6.2 | <0.001 |

Values are mean±SD. Differences between diet groups were determined using parametric unpaired two sample t-test. Males and female were analyzed separately with statistical significance set as $p < 0.05$ . ND, not detectable; NC, not calculable. The lowest detectable limit for both MK4 and deuterium-labeled MK4 was 1pmol/g.

## DISCUSSION

Our findings did not support our hypothesis that dietary supplementation with MK9 will positively influence bone tissue quality and density in mice. In both sexes of mice, femoral tissue strength, section modulus, maximum bending moment, femur length, and bone mineral density did not differ between those fed 0.06 mg MK9/kg diet and those fed 2.1 mg MK9/kg diet. In male mice, moment of inertia and cross-sectional area also did not differ between the diet groups. In female mice, the moment of inertia and cross-sectional area were higher in those receiving 0.06 mg MK9 /kg diet compared to those receiving 2.1 mg MK9 /kg diet. These findings further contradict our hypothesis that higher dietary MK9 intake improves skeletal health.

MK4 and MK9 were found in liver tissues of both sexes. MK9 detected in liver reflects the absorption of vitamin K form supplemented in diet, aligning with a prior study that demonstrated that orally supplemented vitamin K forms accumulated in liver (9). MK4 was the only form of vitamin K detected in bone tissue, with significantly higher concentrations observed in mice receiving the higher dose of MK9. These findings suggest that bone MK4 concentrations respond to dietary MK9 intakes, consistent with what has been reported in other tissues. It was recently demonstrated using stable isotopes that multiple dietary vitamin K forms, including MK9, convert to MK4 in certain tissues (9). However, bone was not measured in that experiment (9). Here we used stable isotopes to evaluate the conversion of dietary MK9 to MK4 in skeletal tissue, and found 63 to 67 % of the MK4 in bone was derived from the labeled MK9 - supporting our hypothesis that dietary MK9 serves as a precursor to MK4 in bone tissue. However, the MK4 concentrations in bone tissue were, for the most part, not correlated with bone tissue quality or density. Moreover, in female mice, higher bone MK4 correlated with a smaller femoral cross-sectional area and a lower moment of inertia (although statistical significance was borderline). Collectively these findings challenge the premise that higher MK4 in bone is beneficial to skeletal health, and the importance of dietary vitamin K conversion to MK4 in bone is unclear.

Results of previous studies focused on MKs and bone health have been inconsistent. In ovariectomized rats, bone strength and BMD were not affected by six weeks of dietary supplementation with 147 mg MK4/ kg diet or 201 mg MK7 /kg diet (equimolar doses) (29). (Like MK9, MK7 is also a bacterially produced form of vitamin K.) In aged female rats, bone formation and histomorphometry did not differ between those fed a diet containing 500 mg MK4/kg diet and those fed a control diet for 98 days (30). However, bone resorption decreased, and bone formation increased in ovariectomized mice fed diets supplemented with 20 and 40 mg MK4/kg body weight (31).

A meta-analysis of randomized clinical trials conducted primarily in post-menopausal women and older adults with osteoporosis found little evidence that vitamin K supplementation has beneficial effects on BMD and inconclusive evidence with respect to fracture risk (13). Most of the included trials supplemented with 45 mg/d of MK4, which is a pharmacological dose that has been prescribed as an anti-osteoporotic medication in Japan (32). Evidence is also mixed regarding the protective effects of MK7. In postmenopausal osteopenic women, BMD and bone microarchitecture did not differ between those who received 375 mcg/d MK7 with 800 mg/d calcium and 1500 IU/d vitamin D for 3 years, and those who received calcium and vitamin D without MK7 (33). Supplementation with 360 µg/d MK7 alone for one year also did not affect BMD in healthy women within five years of menopause (34). However, 180 µg/d MK7 supplementation for 3 years reduced bone mineral density loss at the lumbar spine and femoral neck, but not the total hip, in healthy post-menopausal women. Compression strength and impact strength were also favorably affected by MK7 supplementation (35). This study also did not include calcium or vitamin D supplementation – two nutrients that have been repeatedly shown to protect against age-related bone loss (36, 37). Overall, the available evidence regarding the clinical benefit of MKs with respect to bone health is weak and our findings provide further support for no protective role of MK9 in skeletal health.

Our study has several key strengths. We conducted a comprehensive evaluation of skeletal health in a mouse model, assessing bone tissue quality through multiple measurements as well as bone mineral density. The use of isotopically labeled MK9 allowed us to trace the origin of skeletal tissue MK4 back to dietary MK9, providing robust evidence that MK9 provided in the diet was a source of MK4 in skeletal tissue. However, we acknowledge certain limitations. Some mice, most of which were male, did not survive for the 16-week study. There were no signs of overt bleeding, and the cause of death is unknown. However, this is a consistent finding in male mice when fed low doses of vitamin K for prolonged periods of time (38). Multiple MK forms are produced by bacteria, and it is possible that supplementation with other forms of MK would yield different results. However, previous studies found MK7, a form that has been extensively studied, and MK9 similarly convert to MK4 in extra-hepatic tissues (9), there is no known unique biochemical mechanism for individual MK forms, and evidence regarding the skeletal benefit of MK7 is similarly inconclusive (29, 33–35).

In summary - we found dietary MK9 served as a precursor to MK4 in skeletal tissue, but it did not influence bone tissue quality or BMD. These findings do not support a role for vitamin K in improving skeletal health.

## Abbreviations

^2^H_7_MK9: deuterium-labeled MK9
BMD: bone mineral density
CBU: Comparative Biology Unit
DEXA: dual-energy X-ray absorptiometry
HNRCA: Tufts University Human Nutrition Research Center on Aging
LC-MS: liquid chromatography mass spectrometry
MK: menaquinone
PBS: phosphate-buffered saline
PK: phylloquinone
Q-TOF-MS: time-of-flight mass spectrometry

