## Supplementary Materials for "Dietary Menaquinone-9 Supplementation Does Not Influence Bone Tissue Quality or Bone Mineral Density in Mice"

Supplemental materials

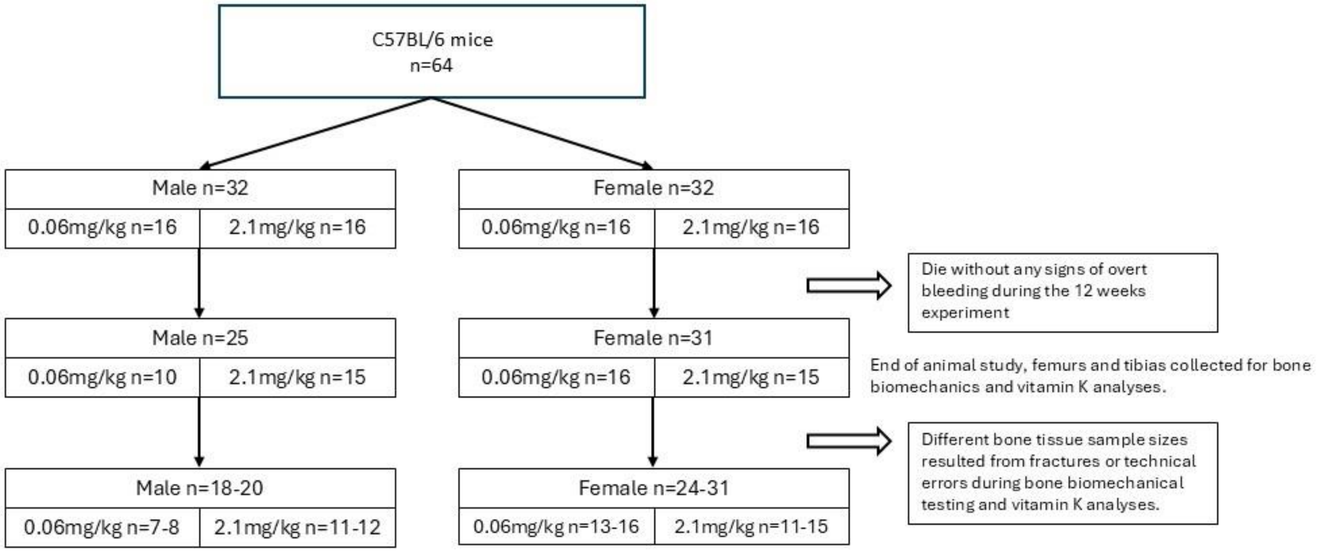


**Figure S1.** Animal Disposition Flowchart. Eight mice died prior to the end of the study (7 males and 1 female) with no signs of overt bleeding. Femurs were unavailable from 7 male and 7 female mice due to bone fractures or technical errors during biomechanical testing. Additionally, tibias from 5 male mice and 0 female mice were lost during vitamin K analyses.

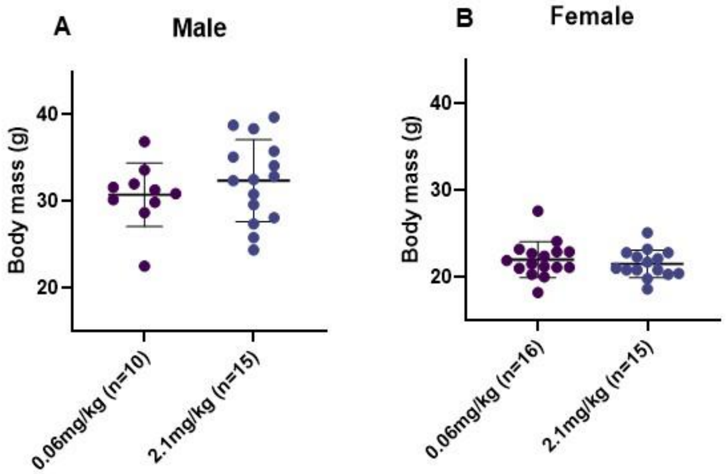


**Figure S2.** Body weight (g) of male **(A)** and female **(B)** mice receiving diets supplemented with MK9. Differences between diet groups were determined using parametric unpaired two sample t-test. Males and female were analyzed separately with statistical significance set as $p<0.05$. Data are presented as mean ±SD. No difference between diet groups in body weight were observed for both sexes (all p values ≥0.345).

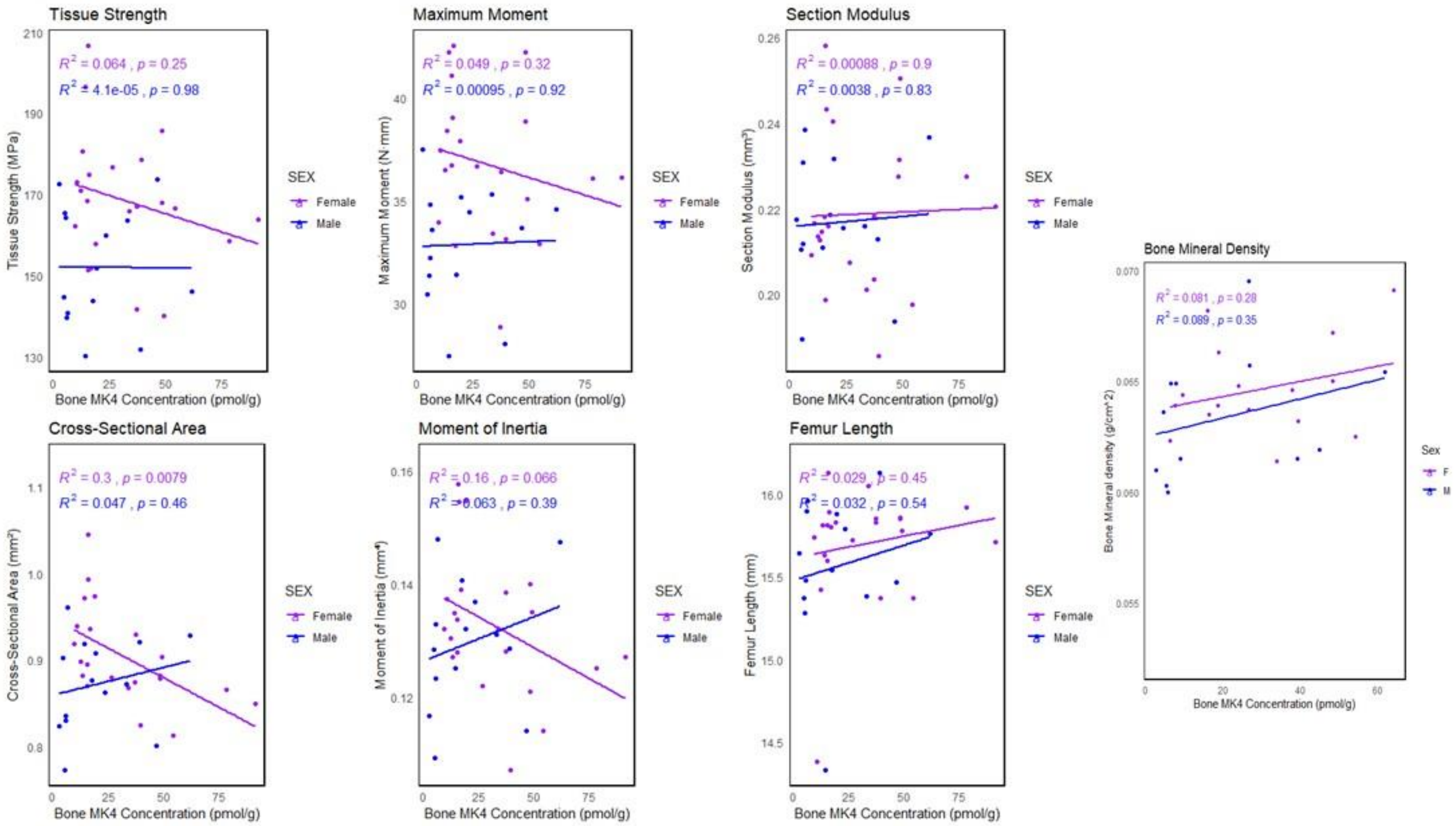


**Figure S3.** Correlations between bone MK4 concentrations (pmol/g) and bone biomechanical and geometry outcomes of male (**blue**) and female (**purple**) mice receiving diets supplemented with MK9. Correlations were assessed using the Pearson correlation coefficient with statistical significance set as $p<0.05$.

**Table S1.** Femoral biomechanics and geometry measurements normalized with body weight of male and female mice receive diets supplemented with MK9.

| | Male | | Female | |
|---|---|---|---|---|
| | 0.06mg/kg (n=7) | 2.1mg/kg (n=11) | 0.06mg/kg (n=13) | 2.1mg/kg (n=11) |
| Tissue strength (Mpa) | 153.3±13.6 | 156±16.3 | 171.8±16.7 | 164.6±14.8 |
| *P values* | 0.758 | | 0.84 | |
| Maximun moment (N*mm) | 33.3±2.4 | 33.7±3.5 | 37.9±3 | 35.4±3.7 |
| *P values* | 0.667 | | 0.309 | |
| Section modulus ($mm^3$) | 0.22±0.02 | 0.22±0.01 | 0.22±0.02 | 0.21±0.02 |
| *P values* | 0.45 | | 0.842 | |
| Moment of inertia ($mm^4$) | 0.13±0.01 | 0.13±0.01 | 0.14±0.01 | 0.13±0.01 |
| *P values* | 0.788 | | 0.01 | |
| Cross sectional area ($mm^2$) | 0.86±0.07 | 0.9±0.04 | 0.95±0.07 | 0.87±0.03 |
| *P values* | 0.937 | | 0.06 | |
| Femur length (mm) | 15.6±0.3 | 15.6±0.5 | 15.6±0.4 | 15.8±0.3 |
| *P values* | 0.458 | | 0.161 | |

The data are presented as original values, mean ± SD. Differences between diet groups were assessed using a parametric unpaired two-sample t-test using **data normalized to body weight**. Males and female were analyzed separately with statistical significant set as $p<0.05$.
